# Structural and Functional Analyses of PolyProline-II helices in Globular Proteins

**DOI:** 10.1101/068098

**Authors:** Prasun Kumar, Manju Bansal

## Abstract

PolyProline-II (PPII) helices are defined as a continuous stretch of a protein chain in which the constituent residues have the backbone torsion angle (φ,ψ) values of (-75°, 145°) and take up extended left handed conformation, lacking any intra-helical hydrogen bonds. They are found to occur very frequently in protein structures with their number exceeding that of π-helices, though it is considerably less than that of α-helices and β-strands. A relatively new procedure, ASSP, for the identification of regular secondary structures using C^α^trace identifies 3597 PPII helices in 3582 protein chains, solved at resolution ≤ 2.5Å. Taking advantage of this significantly expanded database of PPII-helices, we have analyzed the functional and structural roles of PPII helices as well as determined the amino acid propensity within and around them. Though Pro residues are highly preferred, it is not a mandatory condition for the formation of PPII-helices, since ~40% PPII-helices were found to contain no Proline residues. Aromatic amino acids are avoided within this helix, while Gly, Asn and Asp residues are preferred in the proximal flanking regions. These helices range from 3 to 13 residues in length with the average twist and rise being -121.2°±9.2° and 3.0Å±0.1Å respectively. A majority (~72%) of PPII-helices were found to occur in conjunction with α-helices and β-strands, and serve as linkers as well. The analysis of various intra-helical non-bonded interactions revealed frequent presence of C-H…O H-bonds. PPII-helices participate in maintaining the three-dimensional structure of proteins and are important constituents of binding motifs involved in various biological functions.

**ABBREVIATIONS**
MM: main chain-main chain
SM: side chain-main chain
SS: side chain-side chain
SSE: secondary structural elements
H-bond: hydrogen bond
*ASSP*: assignment of secondary structures in proteins
*STRIDE*: Structural identification
std. dev.: standard deviation
*MBA*: Maximum bending angle
*PPII-helix*: PolyProlineII-Helix

## Introduction

Regular secondary structure elements (SSEs) play a vital role in the analysis and understanding of the structure and function of proteins. Among these SSEs, the most abundant α-helices and β-strands were first predicted from theoretical studies (Pauling and Corey, 1951; Pauling et al., 1951) and subsequently confirmed by X-ray diffraction analysis (Blake et al., 1965; Perutz, 1951). It is also a well-known fact that a majority of the helices occurring in globular proteins are the right-handed α-helices with 3_10_-helices being a distant second. Though other types of helices such as π-helices and 2.2_7_ helices are also found in protein structures, their abundance is comparatively very less (Kumar and Bansal, 2015a). The left-handed counterparts of α and 3_10_-helices are rarely found (Hung et al., 1998; Novotny and Kleywegt, 2005; Zawadzke and Berg, 1993). However, PPII-helix is an important class of left-handed helices, which contain about three residues per turn (n=−3) and rise per residue being is ~3.2 Å. PPII-helices are extended structures and the backbone amino as well as carbonyl groups of the constituent residues point away from the helical axis leading to the lack of any intra-helical mainchain-mainchain (MM) N-H…O H-bonds. The residues adopt an average backbone dihedral angles (φ,ψ) of (-75°,+145°) that overlap well with that of the allowed conformational space for Pro residues. Hence, Proline becomes an obvious choice as a constituent residue and Pro-rich regions are often observed to take PPII-helix in protein structures. However, it has been shown that the PPII-helices are also found in the regions of protein or polypeptide that lack significant presence or are completely devoid of Proline residues (Adzhubei et al., 2013). Soman and Ramakrishnan in 1983 (Soman and Ramakrishnan, 1983) reported the presence of PPII-helix in bacteriochlorophyll a-protein and have been since found to occur very frequently in peptide (Makarov et al., 1992) and protein structures (Adzhubei et al., 1987; Stapley and Creamer, 1999). Even the analyses of backbone conformations of individual residues in protein structures deposited in Protein Data Bank (PDB) (Berman et al., 2000) had shown that the PPII and β-strand conformation have comparable occurrences (Adzhubei et al., 1987). Spectroscopic methods have also shown that PPII-helix conformation is a major backbone conformation in unstructured or unordered proteins (Krimm and Tiffany, 1974; Shi et al., 2002). Moreover, it has also been shown that the switch between PPII-helix to either of right-handed α-helix or β-strands can be enabled at the residue level by changing only one dihedral angle (Adzhubei and Sternberg, 1993).

Structural (Beck and Brodsky, 1998; Kieliszewski and Lamport, 1994; Wu et al., 2001) and functional (Chellgren and Creamer, 2004; Kay et al., 2000; Siligardi and Drake, 1995; Stapley and Creamer, 1999) roles of PPII-helices have been discussed in various reports (Williamson, 1994). For example, PPII-helices are shown to play an important role in protein-protein (Kay et al., 2000; Siligardi and Drake, 1995) as well as protein-nucleic acid (Hicks and Hsu, 2004) interactions. The growing number of protein structures with functional and structural roles mediated by PPII-helices has attracted the development of different algorithms like PROSS (Srinivasan and Rose, 1999), XTLSSTR (King and Johnson, 1999), SEGNO (Cubellis et al., 2005), DSSP-PPII (Mansiaux et al., 2011) and ASSP (Kumar and Bansal, 2015b). A dedicated database, PolyprOline, for PPII-helix assignment and analysis has also been developed (Chebrek et al., 2014). Theoretical surveys of PPII-helices in proteins have highlighted their importance (Adzhubei et al., 2013; Brown and Zondlo, 2012; Stapley and Creamer, 1999; Vila et al., 2004). Amino acid propensities for each residue to occur in PPII-helix are also analyzed and compared with those in different SSEs like α-helix, 3_10_-helix and β-strands (Berisio et al., 2006). Host-guest propensity values at the residue level have been analyzed (Kelly et al., 2001), but there is high probability of getting a single residue in PPII conformation, rather than a PPII-helix. However, amino acid propensities in a coiled library made by very restricted values of dihedral angles (φ,ψ) for each SSE showed a good correlation with experimentally derived values (Jha et al., 2005).

Though the importance of PPII-helices in different proteins have been highlighted, a detailed and systematic sequence and structural analysis is essential for the understanding of the principles governing their functions and structural importance. In the present study, we have addressed various questions: i) What are the structural and functional roles played by PPII-helices? ii) What are the factors stabilizing these PPII-helices? iii) What is the position-wise residue propensity within and around PPII-helices? iv) How dissimilar are the position-wise residue propensities within and around PPII-helices and isolated strands? v) Do flanking major secondary structures like α-helices and β-strands affect the position-wise amino acid propensity within and around PPII-helices and finally vi) How different are the PPII-helices, which contain Proline or lack Proline residues? We believe that analyses of these features will provide a better understanding of the PPII-helices and their biological roles.

## Methods

### Composition of Dataset

A dataset of 7957 high-resolution, quality-filtered protein chains termed as top8000 was taken from the Richardson lab (http://kinemage.biochem.duke.edu/databases/top8000.php). Protein chains with length 70 residues and sequence identity >25% (Wang and Dunbrack, 2003) were removed, reducing the number to 3582 chains with 837027 residues.

### Calculation of various local geometric parameters

The computation of various local parameters viz. twist, vtor, rise and radius for each step of four C^α^ atoms has been described in earlier publications (Kumar and Bansal, 2012; Kumar and Bansal, 2015b). A window of four C^α^ atoms slides along the length of the protein, one C^α^ atom at a time. The bending angle at a residue ‘i’ is defined as an angle between the two local helix axes of helical turns encompassing residues from (i-3 to i) and (i to i+3). These bending angles help in identifying the geometry of the overall helix.

### Softwares used

Various helices and β-strands were identified using ASSP and STRIDE. The non-bonded interactions were calculated using MolBridge (Kumar et al., 2014) with default cut-off values. Interactions were further divided into three categories, namely i) main-chain…main-chain (MM), ii) main-chain…side-chain (MS) and iii) side-chain…side-chain (SS). Figures were generated using PyMOL (Schrodinger, 2010) and MATLAB (MATLAB, 2010). Solvent accessibility was calculated using NACCESS (Hubbard and Thornton, 1993).

### PPII-Helix assignment and position nomenclature

Various helices were identified using ASSP and the corresponding numbers are listed in Table 1 along with the median and mean length. A total of 3597 PPII-helices were identified. Algorithm for the identification of PPII-helices has been discussed in the supplementary file. Since the majority of PPII-helices were three residues long, only five positions (N', Ncap, N1, Mid, C1, Ccap and C') within and around them were considered (Figure 1). Three positions (N1, Mid and C1) constitute a helix while the remaining four flanking positions are non-PPII-helical. Unless otherwise stated, N2, N3, N4, C2, C3 and C4 should be considered as a part of ‘Mid’. Unlike other positions, ‘Mid’ may contain one or more residues per helix.

**Figure 1:**
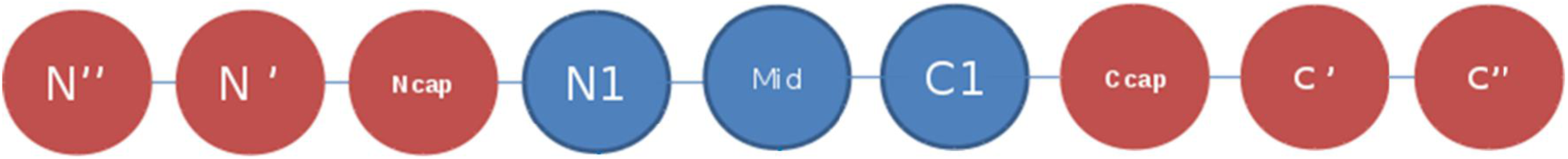
Nomenclature used for various positions within and around PPII-helix. Since majority of PPII-helices were only 3 residues long, 3 helical (in blue) and 6 non-helical positions (in red) were considered for the analyses. However N'-C' were considered for the calculation of position-wise amino acid propensity values. ‘Mid’ position can contain more than one residue.

**Table 1.** Total number of different SSEs as identified by ASSP, DSSP and STRIDE. The mean/ median length corresponds to the SSEs identified by ASSP. Hyphen (-) indicates that the algorithm does not assign the α_L_, 3_10L_, π_L_, PPII fragments. The subscript ‘R’ and ‘L’ defines the handedness of a helix.

| SSEs | ASSP | DSSP | STRIDE | Mean/ Median (Length) |
| --- | --- | --- | --- | --- |
| $\alpha_R$ | 25368 | 24904 | 23102 | 12.6/ 13 |
| $3_{10R}$ | 7594 | 10037 | 10633 | 3.6/ 3 |
| $\pi_R$ | 659 | 857 | 27 | 5.6/ 5 |
| $\alpha_L$ | 17 | - | - | 4.1/ 4 |
| $3_{10L}$ | 36 | - | - | 3.2/ 3 |
| $\pi_L$ | - | - | - | - |
| PPII | 3597 | - | - | 3.5/ 3 |

### Refinement of the dataset consisting of PPII-helices

The comparative analyses of residues in PPII-helices and those in STRIDE or DSSP assigned β-strands have been shown to have small overlap (Kumar and Bansal, 2015b; Mansiaux et al., 2011). To get rid of any ambiguity in the dataset, we compared the residues in PPII-helices to their corresponding assignment by STRIDE. A PPII-helix was considered for further analysis only when more than ‘N/2’ residues of a PPII-helix of length ‘N’ are not identified as a part of β-strand(s) by STRIDE. Representative examples are given in Supplementary Figure SF1. The final dataset consisted of 2879 PPII-helices. Position-wise residue propensity within and around 3516 PPII-helices in the remaining 4375 protein chains were also calculated and compared with those of 2879 PPII-helices in the 3582 protein chains. Both were found to be similar.

### Different categories of PPII-helices

Since the majority of PPII-helices were 3 and 4 residues long, they were divided into 2 sets namely i) PPII^3-4^(PPII-helices of length 3 and 4 residues) and ii) PPII_>4_ (PPII-helices of length > 4 residues). A majority of the PPII-helices were found in conjunction with major SSEs like α-helices or β-strands. Hence, the PPII-helices were divided into four categories based on their occurrence with respect to these major SSEs in a protein chain (Figure 2). If a residue at position C1 of a PPII-helix is the Ncap/N' of the succeeding α-helix or β-strand, the PPII-helix is classified as ‘Nter’, whereas those PPII-helices with N1 being the C'/Ccap of the preceding α-helix or β-strands are termed ‘Cter’. A PPII-helix, sandwiched between two α-helices/β-strands, was categorized as ‘Inter’ (Interspersed), while remaining helices were grouped under ‘Ind’ (Independent) category. Total number of PPII-helices in each category has been listed in Table 2. PPII-helices were also categorized based on the presence of Proline residues. PPII-helices with Pro residues were termed as Pro^+^, while those not containing any Proline were named Pro^-^. A total of 1798 (62.5%) PPII-helices were part of Pro^+^ dataset, while other 1081(37.5%) formed Pro^-^dataset.

**Figure 2:**
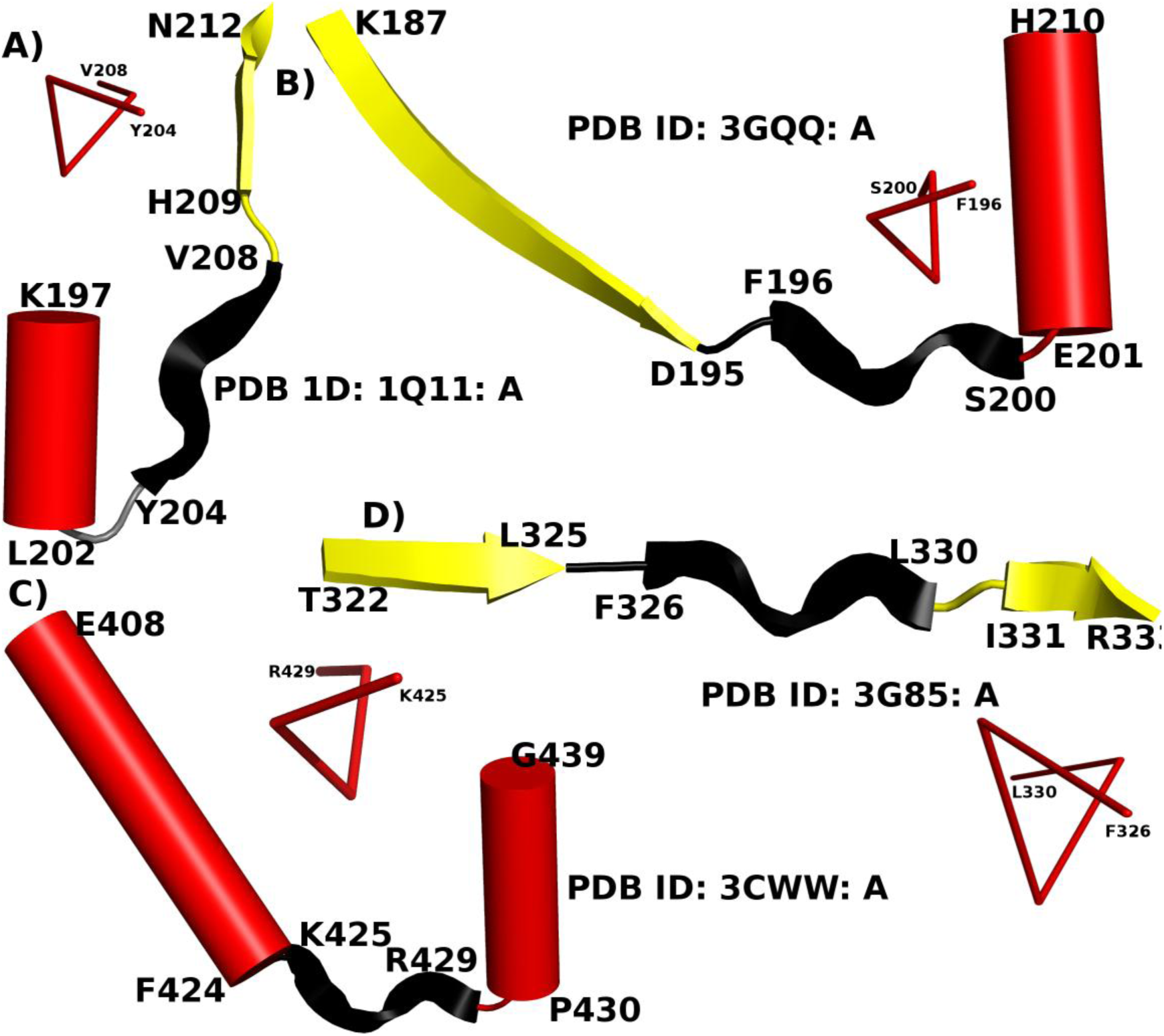
PPII-helices act as a linker between two major SSEs. PPII-helices are colored in black, while α-helix and β-strand are represented as red cylinder and yellow ribbon respectively. The near 3-fold symmetry of PPII-helix region is also shown schematically. The PDB ID and the termini residues of each SSE are given along with each representative example.

**Table 2.** Distribution of PPII-helices in protein structure w.r.t. the flanking major SSEs (α-helix and β-strands). ‘S’ indicates β-strands, while ‘P’ and ‘H’ represent PPII- and α-helix respectively. π- and 3_10_-helices are not considered in the α-helical class.

| Motif | Number (%) | Motif | Number (%) |
| --- | --- | --- | --- |
| SPH | 63 (2.2) | HPS | 61 (2.1) |
| SPS | 69 (2.4) | HPH | 51 (1.8) |
| SP | 650 (22.6) | HP | 330 (11.5) |
| PS | 556 (19.3) | PH | 531 (18.4) |

### Identification of isolated strands

Isolated strands are not part of any β-sheet and like PPII-helices, they also lack any intra MM N-H…O H-bond patterns as the backbone N-H and C=O groups of the constituent residuespoint away from the helix axis. ASSP identified strands of length > 4 residues (β^ASSP^) were compared to those determined by STRIDE. The β^ASSP^, which are not being identified partially or entirely by STRIDE, were considered and subsequently checked for the presence of MM H-bonds also. A total of 393 β^ASSP^ were found to occur in isolation in protein structures.

### Position-wise propensity of residues

Distribution of residues was computed for the 2879 PPII-helices for all 5 positions. Position-wise propensities (P_ij_) for these positions (N'-C') were calculated using the formula used by Kumar and Bansal (Kumar and Bansal, 1998). At a particular position, the preference of residues was determined by ranking them in decreasing order of their propensity values. In order to identify significant changes in the proportion of residues occurring at a particular position, we followed the same methodology as described previously (Kumar and Bansal, 1998).

## Results and Discussion

### PPII-helices commonly occur in globular proteins

ASSP identifies 2879 PPII-helices in 1678 protein chains out of a dataset of 3582 protein chains, with the lactoperoxidase protein from *Bos taurus* (PDB ID: 3NYH: A) containing nine, the maximum number of PPII-helices. Almost every second protein chain was found to have at least one PPII-helix. A total of 9801 (1.2%) residues were involved in the 2879 PPII-helices, which is more than the 3946 (0.5%) residues occurring in 659 π-helices in the same dataset. The length of PPII-helices identified by ASSP ranges from 3 to 13 amino acids, with three residue long PPII-helices constituting almost 72% (2080/2879) of the total. The length and (φ, ψ) distribution of the residues from N1 to C1 positions along with the plot of twist vs. rise and twist vs. radius for 2879 PPII-helices are shown in Figure 3. The mean (μ) and std. dev. (σ) of various parameters are listed in Table 3. The mean twist and rise values were found to be -121.8° and 3.0Å with std. dev. of 8.6° and 0.1Å respectively.

**Figure 3:**
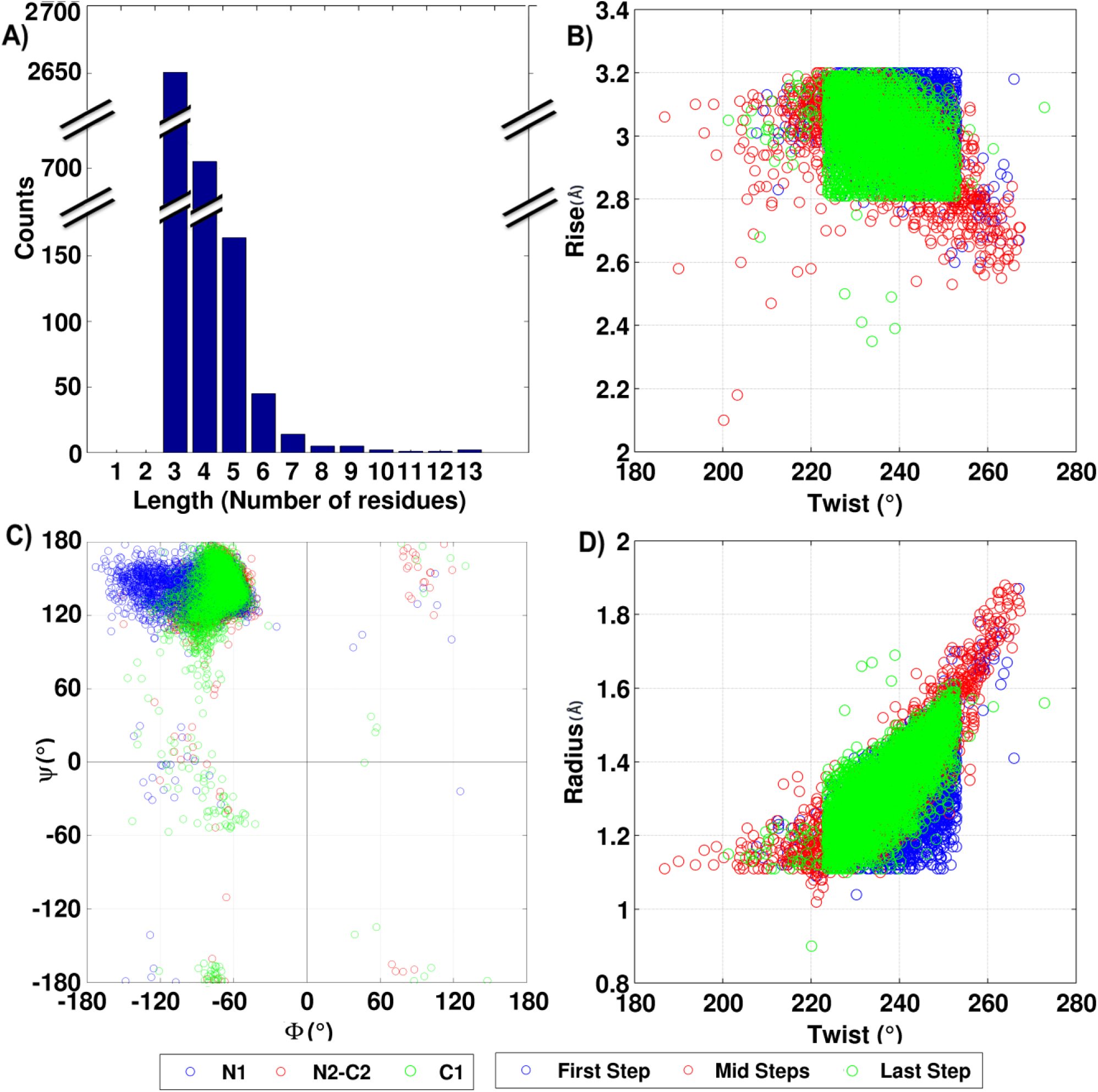
Various parameters characterizing the ASSP identified 2879 PPII-helices: A) length distribution of PPII-helices across the dataset. The longest PPII-helix with 13 amino acid residues (Ala226-Ala238) was identified in peptidase M16 inactive domain (PDB ID: 3GWB: A); B) twist *vs.* rise of 6922 PPII-helical steps; C) distribution of backbone torsion angles (φ, ψ) 9801 residues in PPII-helices; D) twist *vs.* radius of 6922 PPII-helical steps.

**Table 3.** General information and the mean (Std. Dev.) values of different helical parameters in PPII-helices. The corresponding values for β-strands assigned by ASSP are also listed for comparison.

| Parameters | Values |  |
| --- | --- | --- |
| Number of PPII-helices | 2879 |  |
| Number of Steps | 6922 |  |
| Number of Residues | 9801 |  |
| Length (Mean/ Median) | 3.5 / 3 |  |
| | PPII-helix | β-strand |
|  | Mean (Std. Dev.) | Mean (Std. Dev.) |
| Twist ($^{\circ}$) | -121.8 (8.6) | -160.6 (19.2) |
| Rise ($\text{\AA}$) | 3.0 (0.1) | 3.0 (0.4) |
| Radius ($\text{\AA}$) | 1.3 (0.1) | 1.0 (0.2) |
| $\phi$ ($^{\circ}$) | -77.3 (23.2) | -106.1$^{\circ}$ (34.0$^{\circ}$) |
| $\psi$ ($^{\circ}$) | +137.5 (31.5) | +122.4$^{\circ}$ (51.8$^{\circ}$) |

PPII-helices were also checked for their location with respect to the two major SSEs namely, α-helix and β-strand. A total of 531 and 330 PPII-helices were found at the N- and C-terminus of α-helix respectively, while 556 and 650 PPII-helices occurred at the N- and C-terminus of β- strand respectively. Approximately 26% (750) PPII-helices were found to occur independently in protein chains. The presence of PPII-helices interspersed between two major SSEs (Figure 2) suggests that they can often serve as a linker between two SSEs.

### Stabilization of helix termini of PPII-helices

Due to their short length as well as both backbone N-H and C=O groups pointing away from the helical axis, residues in PPII-helices are not involved in intra-helical MM N-H…O H-bonds. These free N-H and C=O groups are often found to form non-bonded interactions with the side chains of flanking non-helical residues, giving rise to characteristic motifs with specific H-bonds and/or (φ-ψ) patterns. Apart from this, the exposed carbonyl or amino groups often extend into the solvent and hence get stabilized by forming a regular network of H-bonds (Kelly et al., 2001; Sreerama and Woody, 1999). The presence of PPII-helix conformation in a seven residue long oligopeptide in a solvent suggests that apart from the electrostatic forces, interactions with the solvent also play a significant role in their stabilization (Rucker and Creamer, 2002). In this article we have discussed C-H…N, C-H…O and N-H…O interactions involving the constituent and flanking residues of the PPII-helices.

The importance of C-H…N non-bonded interactions in determining the crystal structures (Mazik et al., 2000) or molecular packing and conformation (Berkovitch-Yellin and Leiserowitz, 1984; Pickering et al., 2005) of small molecules has been reported, but their role in macromolecules has not been studied. We find that the a total of 2244 C^β^ atoms of (i-1)^th^ residue form C-H…N non-bonded interactions with the backbone amino group of the i^th^ residue (Figure 4A and B). Whereas, 1335 C^γ^ of (i+2)^th^ residues were found to be involved in C-H…O H-bonds with the i^th^ residue. Average donor (D)-Acceptor (A) distances for C-H…N and C-H…O interactions were found to be 3.2Å and 3.7Å respectively, whereas D-H…A angles were found to be 95.6° and 161.3° for C-H…N and C-H…O interactions respectively.

**Figure 4:**
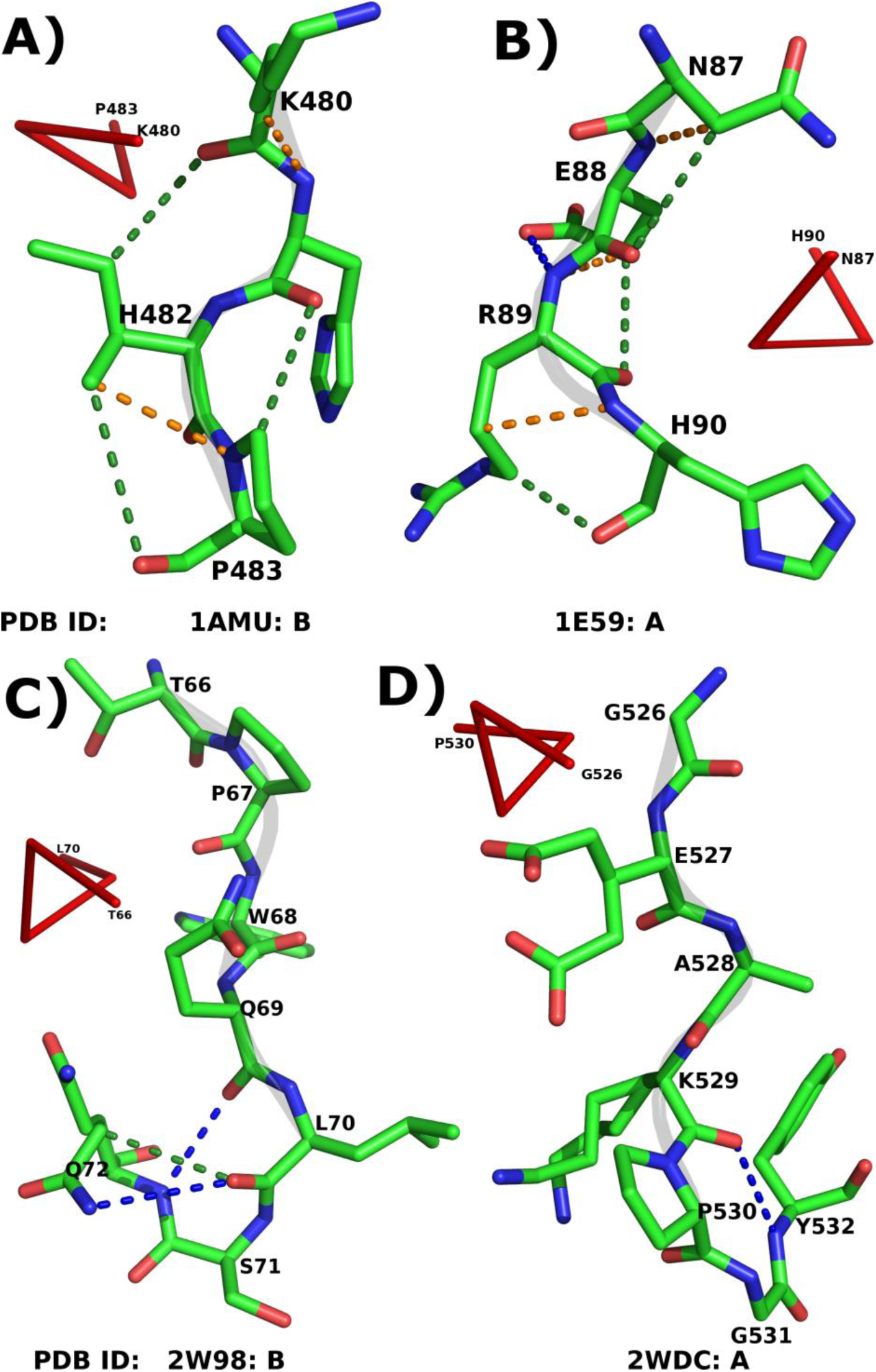
Role of various non-bonded interactions in PPII-helix. Panels A and B illustrate the intra-helical C-H…N (orange dotted lines) and C-H…O (green dotted lines) H-bonds. Panels C and D show the MM N-H…O H-bonds (blue dotted lines) along with C-H…O (if found) at the C-terminus of the PPII-helix. The PDB ID and the termini residues of PPII-helices are given along with each representative example.

The importance of capping motifs in stabilizing the C-terminus of various right handed helices has been studied in detail (Kumar and Bansal, 2015a; Kumar and Bansal, 1998). However, the capping of the PPII-helices is not analyzed to the best of our knowledge. Quite often, the C=O group of C2 residues is capped by the neighboring residues primarily at Ccap and C' positions. At the same time, C1 is shielded by the residues at C' and C''. A total of 230 residues at C2 and 683 residues at C1 were found to be forming MM NH_(i+3)_→O_i_ N-H…O H-bond with flanking non-helical C' and C'' positions respectively (Table 4), suggesting a possibility of β-turns. In order to validate, residues from C2-C' (first case) and C1-C'' (second case) showing NH_(i+3)_→O _i_MM N-H…O H-bond were considered. Turns are characterized by a specific combination of (φ,ψ) values for the 2^nd^ and 3^rd^ constituent residues. Hence, C1 and Ccap have been taken into account in first case, while Ccap and C' in the second case (Supplementary Figure SF2). The average (φ,ψ) of non-Pro and non-Gly residues at Ccap and C' in 2^nd^ case was found to be (-56.6°,-18.2°) and (-64.2˚,-24.7˚) respectively. The mean values for both the positions are similar to the (φ,ψ) values required to form type-III β-turn. Hence, it confirms the role of these turns in capping of the C-terminus of PPII-helices (Figure 4C-D). Additionally, the involvement of C_2_ and C_1_ residues in SM N-H…O H-bonds was also investigated (Table 4). C' and C'' residues were found to be forming the most number of SM N-H…O H-bonds with C_2_ and C_1_ respectively. We conclude that these non-bonded interactions at the C-terminus shield the exposed carbonyl group of the residues and hence stabilize the PPII-helix. However, at the N-terminus of PPII-helix, no such motif was observed and residues were found to be forming H-bonds with the spatially nearby residues or solvent.

**Table 4.** MM (1^st^ row for each position) and SM (2^nd^ row for each position) N-H…O H-bonds formed by the residues at the helix C-termini. Position ‘i’ represents the C2 and C1.MM N-H…O H-bonds were not observed between ‘i’ and ‘i+1’ and hence indicated by Hyphen (–). Similarly for C1, the position ‘i+3’ becomes C''.

| Position | Number | Avg, Dist. D-A ($\text{\AA}$) | Avg. Angle D-H-A ($^{\circ}$) | Number | Avg, Dist. D-A ($\text{\AA}$) | Avg. Angle D-H-A ($^{\circ}$) |
| --- | --- | --- | --- | --- | --- | --- |
|  | C2 (i) |  |  | C1 (i) |  |  |
| (i+1) | - | - | - | - | - | - |
|  | 25 | 2.9 (0.3) | 123.0 (20.7) | 46 | 3.0 (0.3) | 119.3 (22.7) |
| (i+2) | 82 | 3.2 (0.2) | 109.6 (13.0) | 41 | 3.0 (0.2) | 120.1 (16.1) |
|  | 66 | 3.1 (0.2) | 124.9 (23.4) | 318 | 3.1 (0.2) | 142.8 (22.9) |
| (i+3) | 230 | 3.1 (0.2) | 154.0 (13.0) | 683 | 3.1 (0.2) | 139.0 (22.4) |
|  | 80 | 3.1 (0.3) | 154.2 (14.1) | 269 | 3.1 (0.2) | 143.8 (22.0) |

### Functional and structural roles played by PPII-helices in proteins

PPII-helices either directly or indirectly play a major role in facilitating the biological functions and help in maintaining the three-dimensional structure of proteins. The free backbone N-H and C=O groups and extended structure of these helices are essential in facilitating their specific roles. The structural and functional roles of PPII-helices, especially Pro^+^, have been highlighted in various reports (Hicks and Hsu, 2004; Siligardi and Drake, 1995) and reviews (Adzhubei et al., 2013; Williamson, 1994). However, here, we have emphasized on various roles played by the ASSP identified PPII-helices in few representative protein structures.

### Functional importance of PPII-helices

PPII-helices are often found at the surface of the protein and many a time assist protein-protein interactions. For example, a 36-residue polypeptide, avian pancreatic polypeptide from *Meleagris gallopavo* (PDB ID: 1PPT) shows hormonal properties and consists of an α-helix as well as a PPII-helix (Blundell et al., 1981). Both α-helix and PPII-helix run anti-parallel to each other and mainly consist of hydrophobic residues. Two monomers of the protein interact to each other to give a dimer with a hydrophobic core. The formation of hydrophobic core in this molecule is facilitated by the packing together of nonpolar groups, which are constituent residues of PPII-helix and α-helix (Figure 5A).

**Figure 5:**
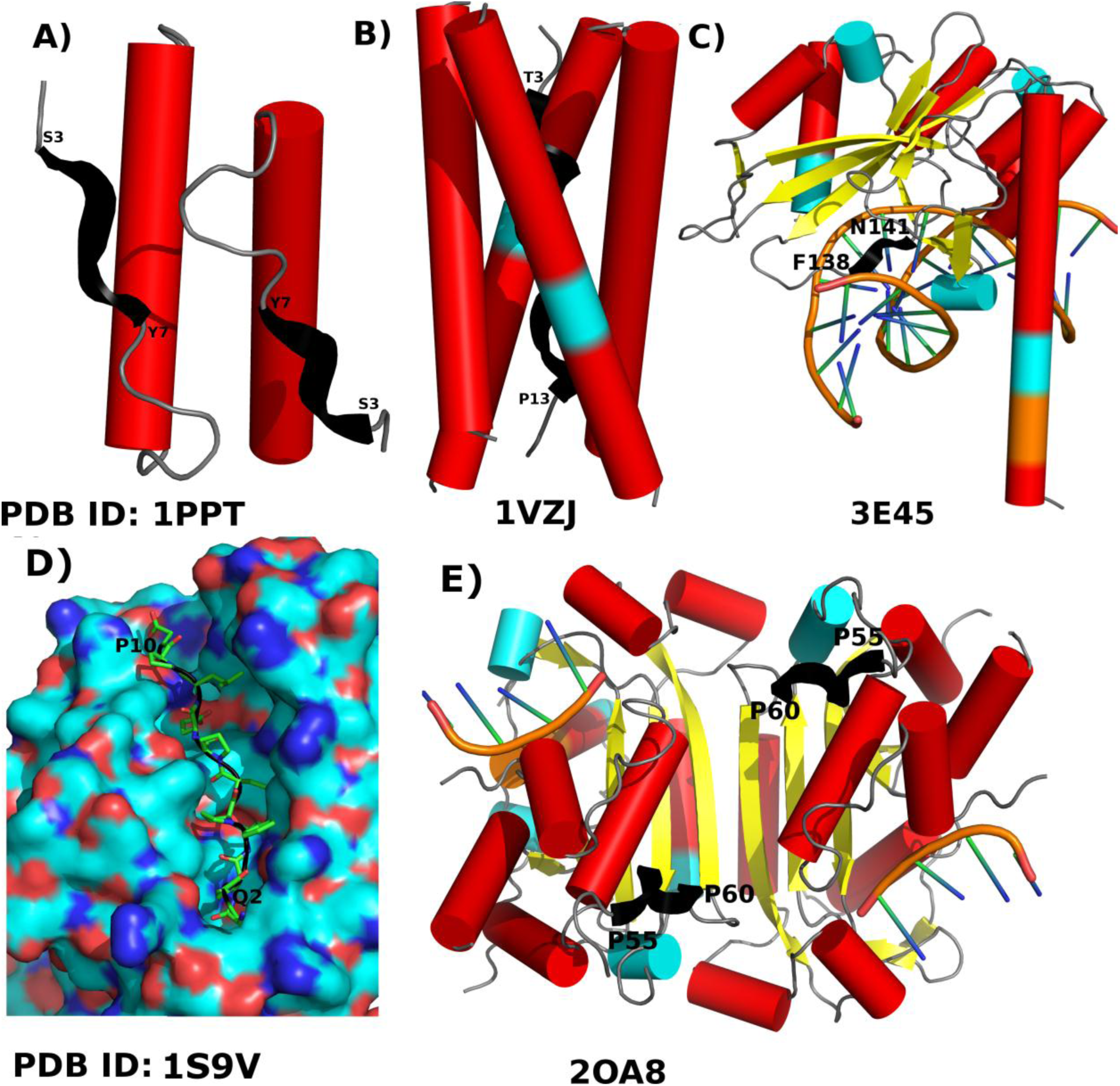
Diverse functional roles of PPII-helices. A) PPII-helix (Ser3-Tyr7) facilitates protein-protein interaction to form a hydrophobic core; B) stabilization of a four helix bundle (PPII-helix: Thr3-Pro13); C) part of a motif involved in Protein-DNA interaction (PPII-helix: Phe138-Asn141); D) PPII-helix is a part of epitope (Pro3-Gln13) and E) PPII-helix (Pro55-Pro60) facilitates the binding of different other proteins. The PDB ID and the termini residues of PPII-helices are given along with each representative example. PPII-helices are colored in black, while α-helix and α-strand are represented as red cylinder and yellow ribbon respectively. 310 and π-helical segments are represented by cyan and orange color cylinders respectively.

PPII-helix also facilitates the formation of tetramerization domain of acetylcholinesterase (AchE) by allowing the four-fold interaction of a WWW motif (PDB ID: 1VZJ) (Dvir et al., 2004). AChE enzymes hydrolyze the acetylcholine and hence terminate the signal transmission at cholinergic synapses. Functional localization of AChE in vertebrate muscle and brain is facilitated by the interaction of the tryptophan amphiphilic tetramerization (WAT) sequence, at the C‐terminus of its major splice variant (T), with a Pro‐rich attachment domain (PRAD; Chain: D), of the anchoring proteins, collagenous (ColQ) and Pro‐rich membrane anchor (Dvir et al., 2004). ASSP assigns PPII-helical conformation to the PRAD (Thr3-Pro13), (Figure 5B), as suggested earlier (Dvir et al., 2004). The main-chain C=O of the residues from PPII-helices were found to form H-bond interactions with the side-chain of the constituent residues of the WAT coiled coils and hence contribute to the overall stability of the WAT/PRAD complex. It has also been reported that the mutation in the PRAD domain disrupts the crucial WAT–WAT and WAT–PRAD interactions (Dvir et al., 2004). Programs like SEGNO and PROSS identified the part of PRAD as PPII-helix, while XTLSSTR was failed to do so.

PPII-helices are even observed to be a part of the structural motif that constitutes DNA-binding regions. Type-II restriction endonuclease HincII enzyme identifies specific short DNA sequence GTYRAC (Y and R indicate any pyrimidine and purine respectively) and perform the endonucleolytic cleavage to give specific double-stranded fragments with terminal 5'-phosphates. The side chain of Gln138 intercalates into the bound DNA just outside of the 6 bp recognition sequence that induces distortions like bending, unwinding, and a shifting of the base planes into the minor groove (Joshi et al., 2006). HincII uses both direct and indirect readout for its activity. The structure of Q138F mutant (PDB ID: 3E45) elucidates the mechanism of indirect readout (Babic et al., 2008). ASSP identifies four residues long PPII-helix from Phe138-Asn141 with π…π stacking between the aromatic ring of Phe138 and Cyt10. Ala139 and Asn141 were also found to form specific non-bonded interactions with the bases Cyt10 (N-H…O) and Ade9 (N-H…N) respectively (Figure 5C). The extended conformation of PPII-helix facilitates specific protein-DNA interactions.

The Pro-rich gliadin peptides naturally adopt a PPII helical conformation and bind to MHC class II molecules and upon deamidation affect the binding as well as increase the immunogenicity (Kim et al., 2004). The protein complex of HLA-DQ2 with an immunogenic epitope from gluten (PDB ID: 1S9V) deciphered the immune-pathogenic basis of celiac disease by studying the interactions between them (Kim et al., 2004). Nine residues long PPII-helix (Gln2-Pro10) binds to the peptide binding groove at the N-terminal domains of DQ2 (Figure 5D). Residues Gln6, Glu8, and Leu9 were found to occupy the P4, P6 and P7 pockets of DQ2 respectively (Kim et al., 2004). The constituent residues, especially Glu8, were found to be involved in various non-bonded interactions. The conformation of the peptide along with the polar nature of component residues facilitates the interaction and hence makes this epitope an excellent ligand for HLA-DQ2.

The structure of TREX1 enzyme (PDB ID: 2OA8) with 3'→5' exonuclease activity reveals an 8-amino acid PPII-helix, suggesting a mechanism for interactions with other protein complexes (De Silva et al., 2007). ASSP assigns PPII-helix to 6 residues (Pro55-Pro60) long Pro-rich segment with a 3-fold symmetry (Figure 5E). PPII-helices in TREX1 dimer have been illustrated to function as interaction motifs with other proteins containing SH3, WW or EVH1 domains (Zarrinpar et al., 2003). The distance between PPII-helix of each monomer of a dimer was found to be 20Å and positioned on the same face of the dimer. Hence, the positioning of Pro-rich PPII-helix plays a key role in protein-protein interaction for the TREX1 protein (De Silva et al., 2007).

### Structural importance of PPII-helices

PPII-helices can act as a linker between two structural or functional domains. For example, parvalbumin molecule belongs to EF-hand_7 (PF13499) family (Punta et al., 2012) and has helix-loop-helix topology. In a crystal structure of rat α-parvalbumin (PDB ID: 1RWY: B) (Bottoms et al., 2004), ASSP identified PPII-helix (Ala74-Ser78) that is a part of linker connecting the two functional domains namely CD and EF (Figure 6A). These domains show Ca^2+^ binding activity and they have been reported to be related by a twofold symmetry axis (Kretsinger and Nockolds, 1973). The extended conformation of the PPII-helix enables the residue Arg75 to form a salt-bridge with Glu81 that plays a vital role in stabilizing the loop region joining two helical segments.

**Figure 6:**
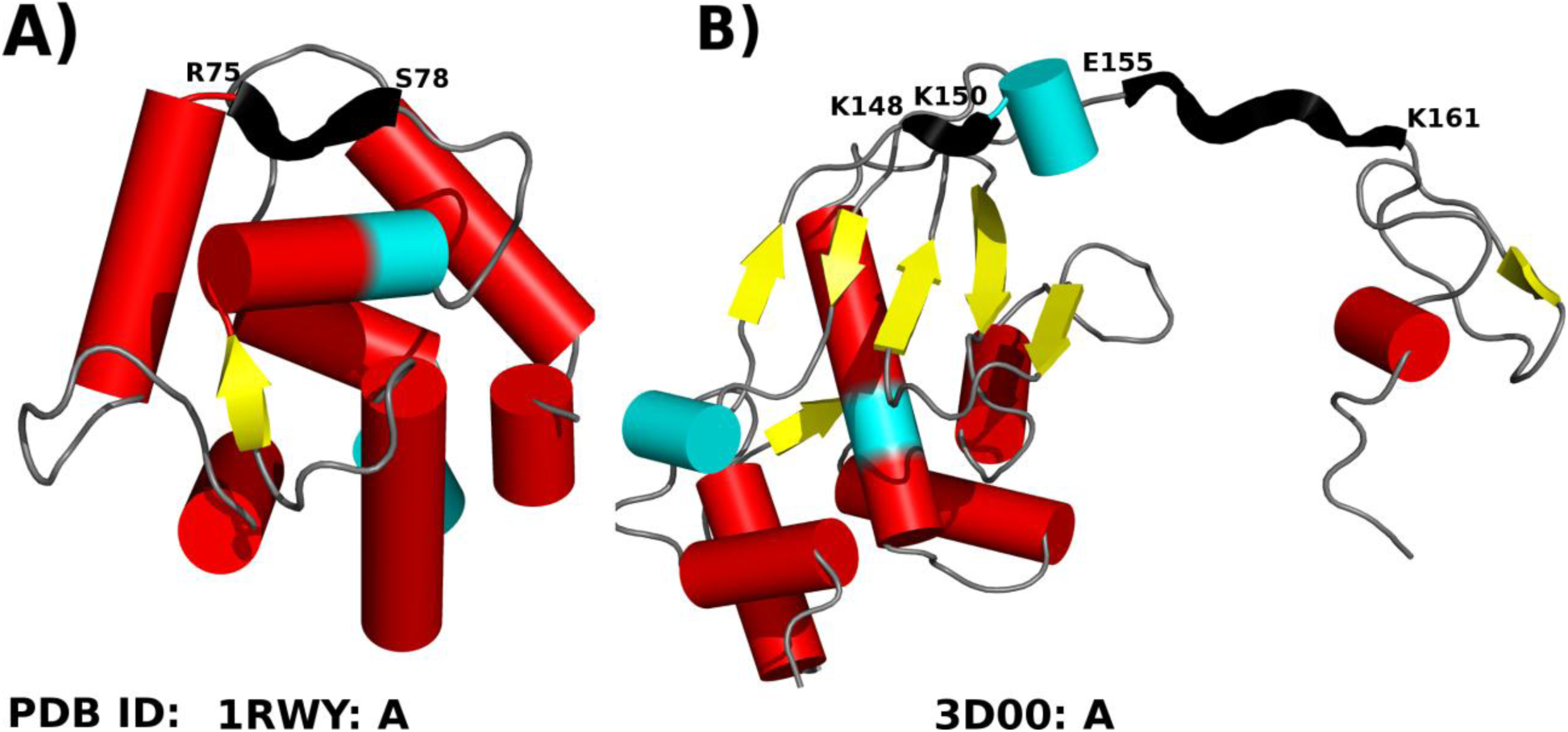
Structural role of PPII-helix. PPII-helix acts as a linker between A) two functional domains and B) two structural domains. The PDB ID and the termini residues of PPII-helices are given along with each representative example. PPII-helices are colored in black, while α-helix and β-strands are represented as red cylinder and yellow ribbon respectively. 3_10_-helical segments are represented by cyan color cylinders.

Formylmethanofuran dehydrogenases are multi-subunit enzymes that catalyze the first step in the formation of CH_4_ from CO_2_ in methanogenic and sulfate-reducing microorganisms (Thauer et al., 2008). They also contain tungsten or molybdenum as well as iron–sulfur clusters. Crystal structure of tungsten formylmethanofuran dehydrogenase subunit-e (PDB ID: 3D00:A; fmde; Pfam ID: PF02663)-like protein from *syntrophus aciditrophicus* have been reported to have PPII-helix (Axelrod et al., 2010). The C-terminal domain is anchored to the N-terminal domain with the help of an eleven residues linker and ASSP assigns PPII-helix (Gln155-Lys161) to a part of it (Figure 6B). The extended structure of the PPII-helix allows the two domains to be separated optimally to form a dimer with swapped-domains. To our surprise, in both the cases, the PPII-helices were found to contain only non-Pro residues.

PPII-helices have also been shown to provide local order, flexibility or chain hydration. It is also one of the predominant conformational states. Spectroscopic data analyses (Shi et al., 2006), CD (Whittington et al., 2005) and solid-state NMR analyses (Hu et al., 2009) have indicated that the PPII-helices maintain local order in largely unfolded proteins or peptides. Additionally, PPII-helices also play a vital role in maintaining a 3D structure of the proteins associated with conformational diseases (Blanch et al., 2000; Blanch et al., 2004; Syme et al., 2002) such as Alzheimers.

PPII-helices are also shown to be an integral part of snow flea antifreeze protein (PDB ID: 3BOI) (Pentelute et al., 2008). The structure consists of 6 PPII-helices that are stacked in two sets of three and form a compact brick-like structure. In another example, the Ala-rich domain and Pro-rich segment in A3VP1 of AgI/II of *Streptococcus mutans* (PDB ID: 3IPK) adopt an extended α-helix and PPII-helix respectively (Larson et al., 2010). Both α-helix and PPII-helix interlock with each other to form a highly extended stalk-like structure (Larson et al., 2010) that can extend over 50 nm in length.

### Residue preferences in PPII-helices depend on the presence of Pro

The position-wise propensity of 20 residues to occur at each of the five positions of PPII-helices of different lengths (PPII _3-4_and PPII_>4_; defined in Methods) were calculated and analyzed (Figure 7).

**Figure 7:**
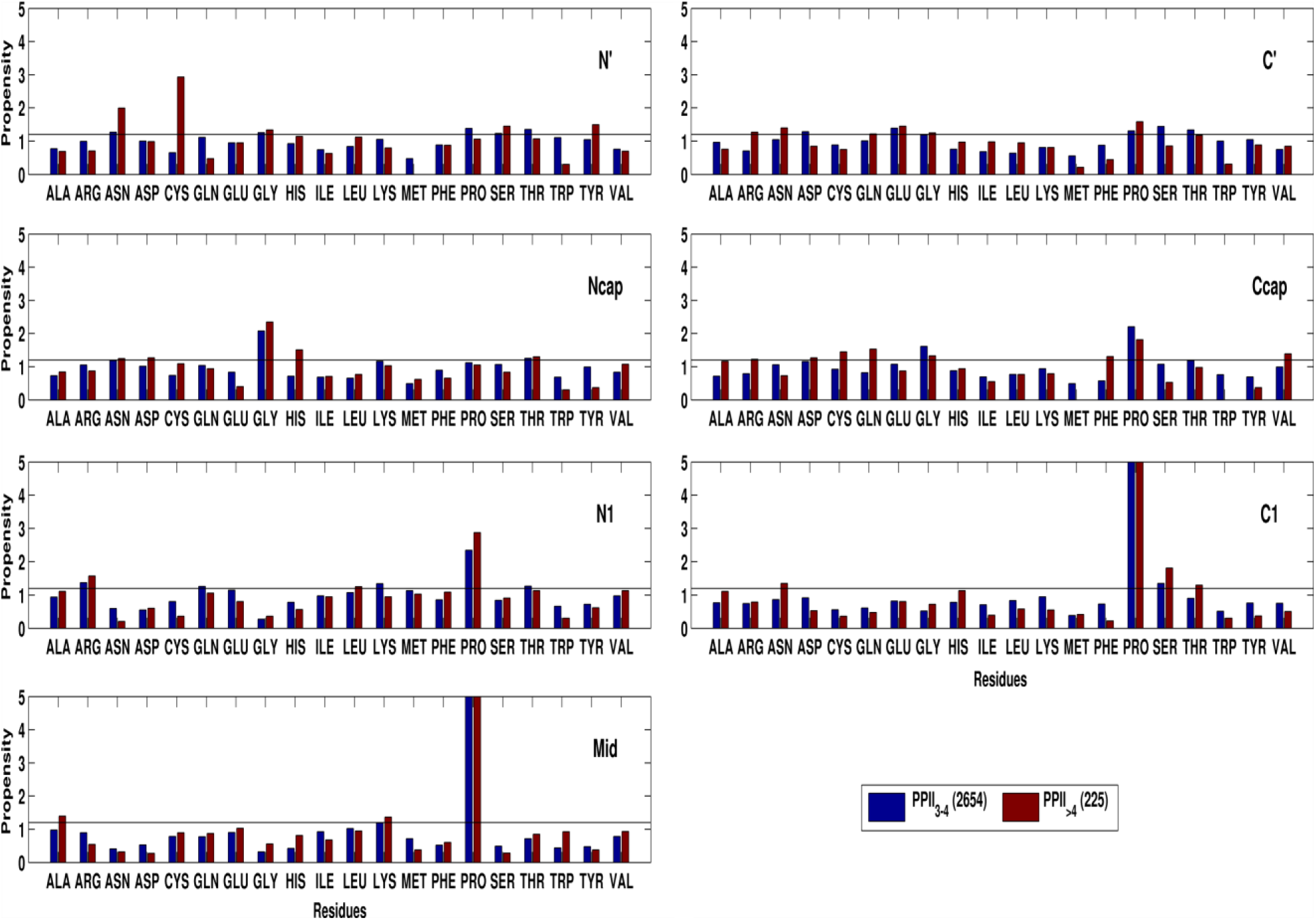
Position-wise amino acid preferences for 20 standard amino acid residues for different length categories of PPII-helices. The horizontal line at 1.2 corresponds to the propensity cutoff value, above which the residue occurrence is considered as significant. The position-wise amino acid propensity of Pro was found to be >5 for both PPII_3-4_ and PPII_>4_ at Mid (6.3, 5.6) and C1 (5.2, 6.2) positions.

PPII_>4_ showed a preference for Cys and Asn at N'. However, at Ncap position, Gly was found to be the most preferred residue for both length dependent categories of PPII-helices. Additionally, PPII_>4_ showed a preference for His also. As expected, our analyses showed a remarkably high preference for Pro residues at all the helical positions (N1-C1) of PPII-helices as suggested earlier (Berisio et al., 2006; Jha et al., 2005; Kelly et al., 2001; Stapley and Creamer, 1999). However, in the case of PPII_>4_, Arg is also preferred at N1, while Ala and Lys are preferred at Mid positions. At C1 position of PPII_>4_ Asn, Ser and Thr were also preferred. Surprisingly, both PPII_3-4_ and PPII_>4_ showed a preference for Gly at Ccap position. Additionally, PPII_>4_ were found to prefer Gln and Val also. Polar residues like Asn and Glu are preferred by both PPII_3-4_ and PPII_>4_ at C' position. In addition to it, PPII_3-4_ showed a preference for Asp and Ser also. Contrary to the previously reported significant correlation between the amino acid propensities for PPII-helices and α-helices (Berisio et al., 2006), results from our analyses suggest that the position-wise amino acid propensity at various helical positions in PPII-helices is different than that of α-helices (Kumar and Bansal, 2015a; Kumar and Bansal, 1998) or π-helices (Kumar and Bansal, 2015a). α-helices have unique preference for amino acids at various positions, while PPII-helices have the overwhelming preference for Proline at almost all the positions. However, π-helices prefer to have hydrophobic or aromatic residues.

Almost 40% of the total PPII-helices were found to contain residues other than Pro (Pro^-^). The longest Pro^-^ in our dataset was found to be 11 residues long (Gly15-Gly25) in antifreeze protein (PDB ID: 3BOI). The structural and functional roles of such Pro^-^ have been discussed in different proteins (Adzhubei et al., 2013). The position-wise propensity of 19 remaining residues in such helices was calculated, analyzed and compared with Pro^+^ and α-helices of length < 9 residues (α_4-8_). It was observed that the higher preference for Pro residues in Pro^+^ has been compensated by the abundance of polar residues in Pro^-^ (Figure 8). Ncap position for both Pro^+^ and Pro-showed a preference for Gly residue; whereas mainly polar residues were preferred in (α_4-8_). However, Asn and Lys are also preferred at Ncap of Pro^-^. At N1, Pro^-^showed a higher preference for Arg, Glu, Lys and Met residues, while α_4-8_ preferred Pro and Trp. At C1, the elevated preference for Pro in Pro^+^ was compensated by higher propensity for Asn, Ser and Thr. However, Leu, Asn and Gln were favored in α_4-8_. Lys, Glu and Ala have higher position-wise amino acid propensity value at Mid position of Pro^-^, while α_4-8_ preferred Leu, Glu and Ala. Both Pro^-^ and Pro^+^, preferred Pro at the Ccap position, while Gly has a significantly high preference in α_4-8_.Surprisingly, at C', both categories of PPII-helix did not show any significant difference. However, Pro with position-wise propensity, P_ij_=1.9 was found to be the most preferred residue at C' of α_4-8_.

**Figure 8:**
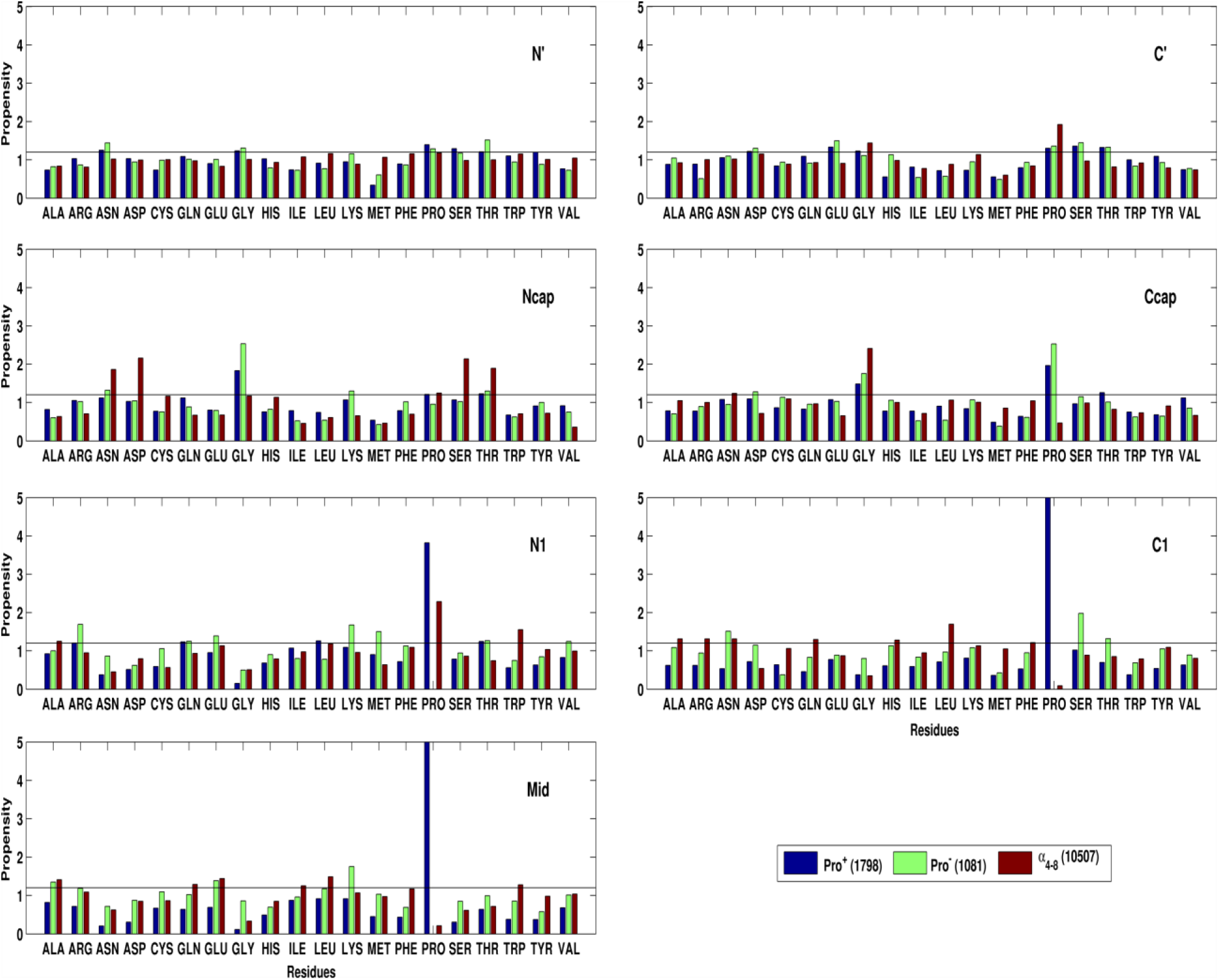
Position-wise amino acid preferences for 20 standard amino acid residues in Pro^+^, Pro^-^ and α_4-8_. The horizontal line at 1.2 corresponds to the propensity cutoff value, above which the residue occurrence is considered as significant. The position-wise amino acid propensity of Pro in Pro^+^ was found to be >5 at Mid (9.4) and C1 (8.4) positions.

The preference for polar residues in PPII-helices led us to investigate their solvent accessibility. A residue was considered solvent exposed when the relative accessibility value is ≥ 7 for all atoms. It was observed that in ~87% (2489/2879) of PPII-helices, more than 50% constituent residues are solvent exposed. A total of 390 PPII-helices were found to be solvent occluded. Even, these 390 PPII helices have a preference for Pro at different helical positions. Additionally, Met and Cys at N1 and Mid respectively have substantial position-wise amino acid propensity values. Surprisingly, at Ncap, the propensity for Pro was not found to be significant. It was compensated by a variety of residues like Gly, Thr, Phe, Asn, Arg and Gln. However, Ccap showed high position-wise propensity values for Gly and Pro.

### Comparison of PPII helices and isolated extended strands

Isolated strands (β^ASSP^) are not part of any β-sheet and lack MM N-H…O H-bonds. Hence, β^ASSP^ is similar to the PPII-helices in terms of absence of any MM N-H…O H-bond pattern. It motivated us to compare the position-wise amino acid propensity within and around β^ASSP^ and PPII-helices. A total of 225 PPII-helices and 393 β^ASSP^ of length > 4 residues were considered (Figure 9). Since both PPII _>4_and β^ASSP^ are of minimum five residues length, five helical (N1, N2, Mid, C2 and C1) as well as four non-helical (N', Ncap, Ccap and C') positions were considered for the comparison of amino acid propensity values. Interestingly both SSEs show a high preference for Proline residues at all the helical positions, but it was invariably found to be higher for PPII_>4_.

**Figure 9:**
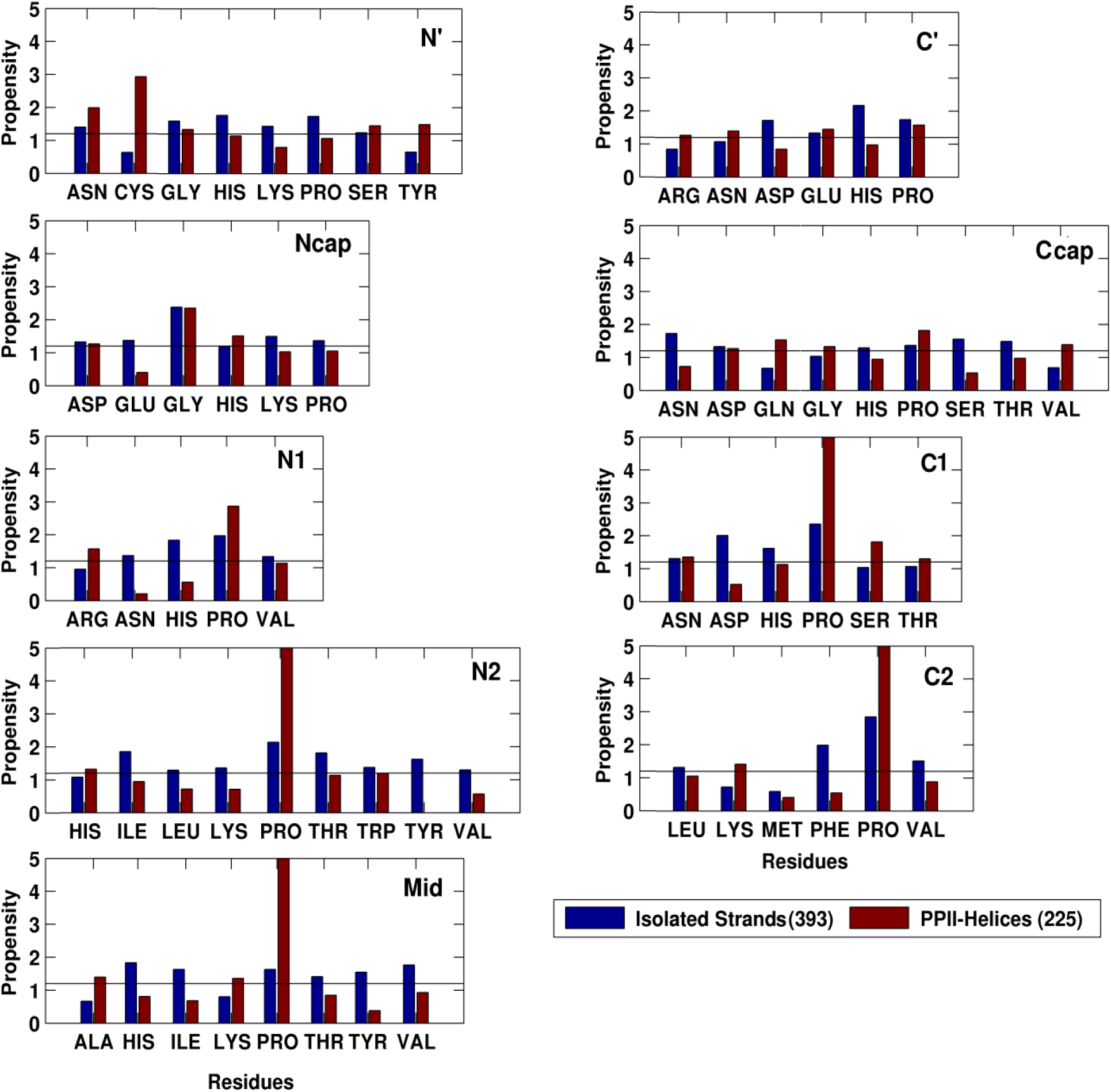
Position-wise amino acid preferences for selected amino acid residues showing significant differences for isolated strands and PPII-helices of length > 4 residues. The horizontal line at 1.2 corresponds to the propensity cutoff value, above which the residue occurrence is considered as significant. The position-wise amino acid propensity values (P_ij_) of Pro at N2, Mid, C2 and C1 in PPII-helices were found to be 6.6, 5.6, 6.2 and 5.7 respectively.

Lys and Pro showed significantly higher propensity at N' for β^ASSP^ compared to Cys in PPII_>4_. However, Gly and Asn are preferred in both. At Ncap both showed the preference for Gly. N1 position in β^ASSP^ preferred Asn and His residues. N2 showed a higher preference for hydrophobic residues (Ile, Leu and Val) and aromatic amino acids (Thr, Tyr and Trp) in β^ASSP^ compared to His in PPII_>4_. At Mid, β-branched residues like Ile and Val are preferred in β^ASSP^ along with aromatic residues Thr and Tyr. Val and Phe are preferred at C2 in β^ASSP^, while Lys in PPII_>4_. Unlike other intra-β^ASSP^ positions, C1 and Ccap showed a preference for polar residues. Asp is preferred at C1, while Asn, Ser and Thr are favored at Ccap. At C', Asp and His were found to be preferred. The preference of polar residues at helical positions N1 and C1 of β^ASSP^can be attributed to the formation of SM H-bonds leading to their stability.

## Conclusions

A total of 2879 PPII-helices were identified by ASSP in a dataset of 3582 protein chains, suggesting that PPII-helices occur quite frequently in proteins. Though Pro residues are preferred in PPII-helices, almost 40% of total helices do not have any Proline. Our analysis suggests that the type III β-turn is a common capping motif at the C-terminus of PPII-helices. PPII-helices in proteins mediate various structural and functional roles. PPII-helices show characteristic preference for specific residues at various helical and flanking non-helical positions. It is different from that of other right-handed helices like α or π. Gly was found to be preferred at both N- and C-terminus of PPII-helix. The higher preference of Pro in Pro^+^helices was compensated by the preference for polar residues in Pro^-^. Though isolated strands and PPII-helices both lack any intra-helical MM N-H…O H-bonds, they differ from each other in their amino acid preference at various positions.

## Supplementary File

### 1.1 ASSP and identification of PPII-helices: Determining cut-off values of local helical parameters

ASSP uses only C^α^ atoms for calculating the local helical parameters (twist, rise, vtor etc.) and identify continuous stretches in a given protein chains (Kumar and Bansal, 2015b). Depending on the cut-off values of the helical parameters, these continuous stretches are further characterized into various secondary structure elements. Here, we have defined the steps followed for the identification of PPII-helices during development of ASSP. The initial estimation of PPII regions in a protein chain is based on criteria described by Stapley and Creamer (Stapley and Creamer, 1999). The cut-off values of various parameters for PPII-helices were determined using the continuous stretches, for which the backbone torsion angles (φ, ψ) satisfy the following criteria: (a)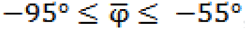, 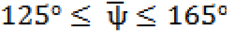; (b) 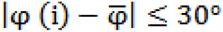, 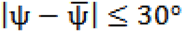 and (c) 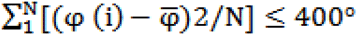, 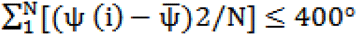. φ (i) and ψ (i) are the backbone torsion angles of i^th^ residue, while are the mean values of backbone torsion angles for a N residues long continuous stretch. In a dataset of 1008 protein folds (used as a training set for ASSP), a total of 254 continuous stretches were found to satisfy the above criteria of which 123 continuous stretches were found to have partial or complete overlap with the STRIDE assigned β-strands and hence not considered for the cut-off calculation. Using the step parameters of 405 steps from remaining 131 continuous stretches, the mean and Std. Dev. values were calculated. Lower and upper range was taken as the mean ± Std. Dev (Supplementary Table ST1). Applying the cut-off values of these local helical parameters to the continuous stretches, a total of 824 PPII-helices were identified (Kumar and Bansal, 2015b).

**ST1:** Mean (Std. Dev.) values obtained by using 405 steps from 131 continuous stretches. lower and upper limit values are defined by considering mean ± Std. Dev.

| Parameters | Mean (Std. Dev.) | Lower Limit | Upper Limit |
| --- | --- | --- | --- |
| Twist (°) | 238.8 (15.2) | 223.6 | 254.0 |
| Rise (Å) | 3.0 (0.2) | 2.8 | 3.2 |
| Vtor (°) | 250.7 (19.5) | 231.2 | 270.2 |
| Radius (Å) | 1.5 (0.4) | 1.1 | 1.9 |

**SF1:**
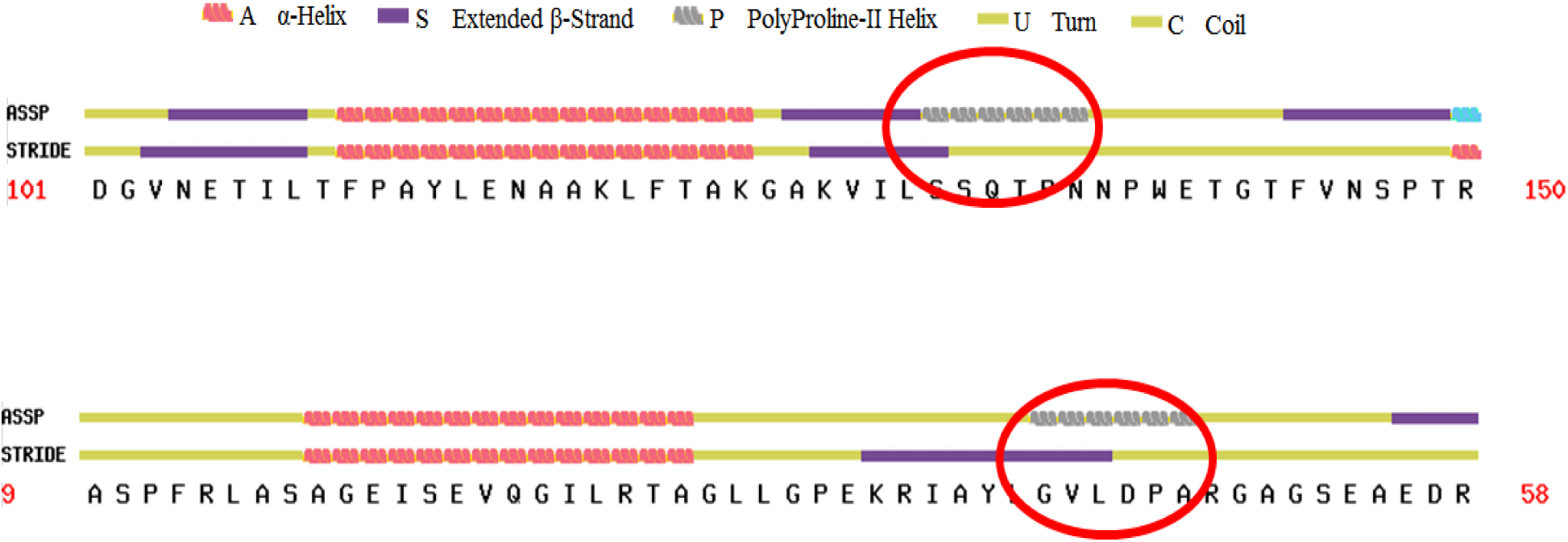
Procedure for refinement of PPII-helices. A PPII-helix will not be considered for analyses, if it has 50% or more overlap with the STRIDE assigned β-strand. Since the overlap is less than 50% in the top panel (PDB ID: 1K7C: A), the PPII-helix will become part of the curated dataset. Whereas, the same is not the case with the PPII-helix in the bottom panel (PDB ID: 2CG1: A).

**SF2:**
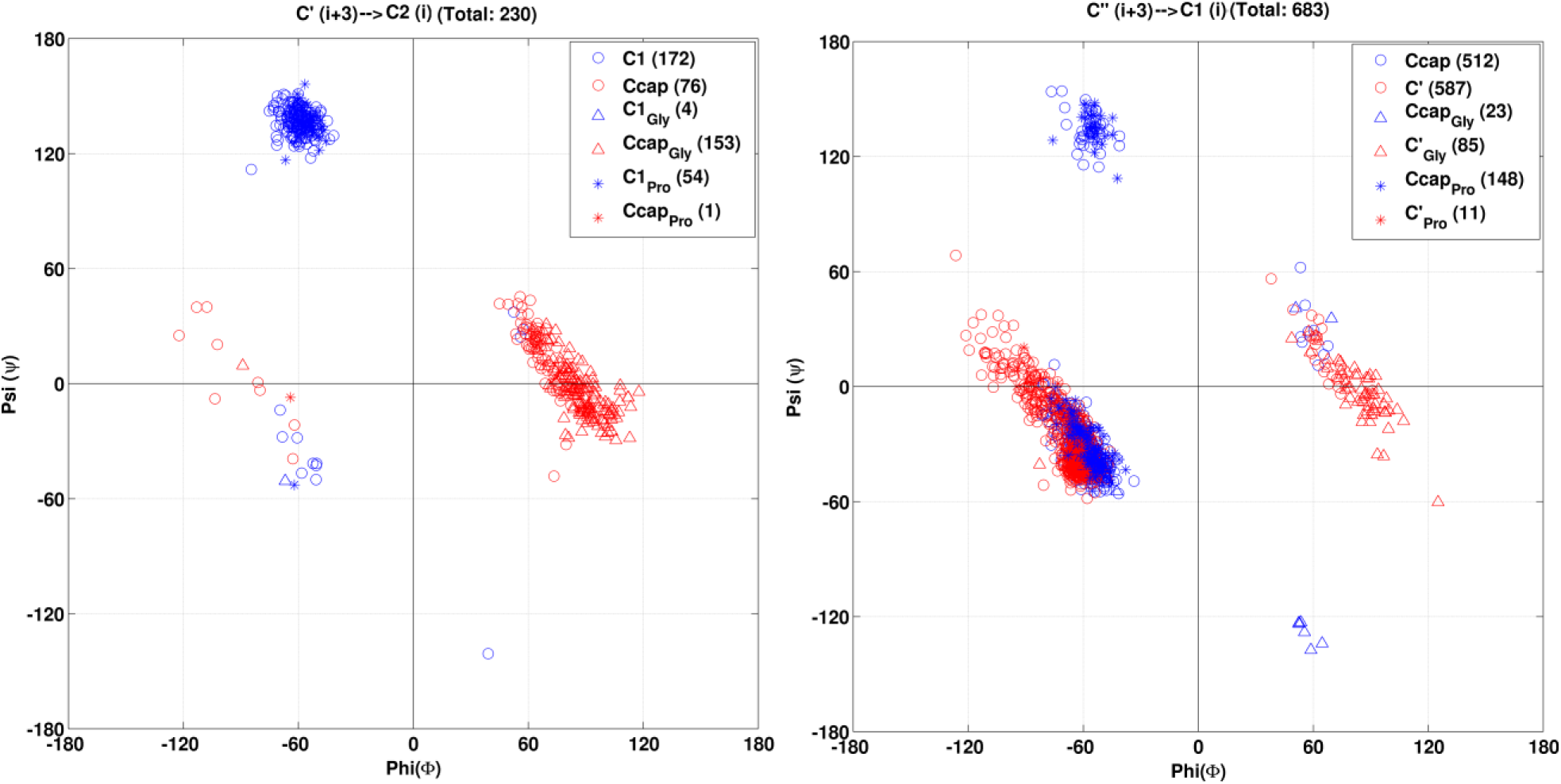
(φ,ψ) distribution of 2^nd^ and 3^rd^ residues of turn formed by (A) C2-C1-Ccap-C' and (B) C1-Ccap-C'-C''. Data points shown in hollow circle are for all non-Pro and non-Gly residues.

## References

Adzhubei, A.A., Sternberg, M.J., 1993. Left-handed polyproline II helices commonly occur in globular roteins. J Mol Biol 229, 472–493.

Adzhubei, A.A., Sternberg, M.J., Makarov, A.A., 2013. Polyproline-II helix in proteins: structure and function. J Mol Biol 425, 2100–2132.

Adzhubei, A.A., Eisenmenger, F., Tumanyan, V.G., Zinke, M., Brodzinski, S., Esipova, N.G., 1987. Third type of secondary structure: noncooperative mobile conformation. Protein Data Bank analysis. Biochem Biophys Res Commun 146, 934–938.

Axelrod, H.L., Das, D., Abdubek, P., Astakhova, T., Bakolitsa, C., Carlton, D., Chen, C., Chiu, H.-J., Clayton, T., Deller, M.C., Duan, L., Ellrott, K., Farr, C.L., Feuerhelm, J., Grant, J.C., Grzechnik, A., Han, G.W., Jaroszewski, L., Jin, K.K., Klock, H.E., Knuth, M.W., Kozbial, P., Krishna, S.S., Kumar, A., Lam, W.W., Marciano, D., McMullan, D., Miller, M.D., Morse, A.T., Nigoghossian, E., Nopakun, A., Okach, L., Puckett, C., Reyes, R., Sefcovic, N., Tien, H.J., Trame, C.B., van den Bedem, H., Weekes, D., Wooten, T., Xu, Q., Hodgson, K.O., Wooley, J., Elsliger, M.-A., Deacon, A.M., Godzik, A., Lesley, S.A., Wilson, I.A., 2010. Structures of three members of Pfam PF02663 (FmdE) implicated in microbial methanogenesis reveal a conserved [alpha]+[beta] core domain and an auxiliary C-terminal treble-clef zinc finger. Acta Crystallographica Section F 66, 1335–1346.

Babic, A.C., Little, E.J., Manohar, V.M., Bitinaite, J., Horton, N.C., 2008. DNA distortion and specificity in a sequence-specific endonuclease. J Mol Biol 383, 186–204.

Beck, K., Brodsky, B., 1998. Supercoiled protein motifs: the collagen triple-helix and the alpha-helical coiled coil. J Struct Biol 122, 17–29.

Berisio, R., Loguercio, S., De Simone, A., Zagari, A., Vitagliano, L., 2006. Polyproline helices in protein structures: A statistical survey. Protein Pept Lett 13, 847–854.

Berkovitch-Yellin, Z., Leiserowitz, L., 1984. The role played by C-H…O and C-H…N interactions in determining molecular packing and conformation. Acta Crystallographica Section B 40, 159–165.

Berman, H.M., Westbrook, J., Feng, Z., Gilliland, G., Bhat, T.N., Weissig, H., Shindyalov, I.N., Bourne, P.E., 2000. The Protein Data Bank. Nucleic Acids Res 28, 235–242.

Blake, C.C., Koenig, D.F., Mair, G.A., North, A.C., Phillips, D.C., Sarma, V.R., 1965. Structure of hen egg-white lysozyme. A three-dimensional Fourier synthesis at 2 Angstrom resolution. Nature 206, 757–761.

Blanch, E.W., Morozova-Roche, L.A., Cochran, D.A., Doig, A.J., Hecht, L., Barron, L.D., 2000. Is polyproline II helix the killer conformation? A Raman optical activity study of the amyloidogenic prefibrillar intermediate of human lysozyme. J Mol Biol 301, 553–563.

Blanch, E.W., Gill, A.C., Rhie, A.G., Hope, J., Hecht, L., Nielsen, K., Barron, L.D., 2004. Raman optical activity demonstrates poly (L-proline) II helix in the N-terminal region of the ovine prion protein: implications for function and misfunction. J Mol Biol 343, 467–476.

Blundell, T., Pitts, J., Tickle, I., Wood, S., Wu, C.-W., 1981. X-ray analysis (1. 4-Å resolution) of avian pancreatic polypeptide: Small globular protein hormone. Proceedings of the National Academy of Sciences 78, 4175–4179.

Bottoms, C.A., Schuermann, J.P., Agah, S., Henzl, M.T., Tanner, J.J., 2004. Crystal structure of ratα‐parvalbumin at 1.05 Å resolution. Protein science 13, 1724–1734.

Brown, A.M., Zondlo, N.J., 2012. A propensity scale for type II polyproline helices (PPII): aromatic amino acids in proline-rich sequences strongly disfavor PPII due to proline–aromatic interactions. Biochemistry 51, 5041–5051.

Chebrek, R., Leonard, S., de Brevern, A.G., Gelly, J.C., 2014. PolyprOnline: polyproline helix II and secondary structure assignment database. Database (Oxford) 2014.

Chellgren, B.W., Creamer, T.P., 2004. Short sequences of non-proline residues can adopt the polyproline II helical conformation. Biochemistry 43, 5864–5869.

Cubellis, M.V., Cailliez, F., Lovell, S.C., 2005. Secondary structure assignment that accurately reflects physical and evolutionary characteristics. BMC Bioinformatics 6, S8.

De Silva, U., Choudhury, S., Bailey, S.L., Harvey, S., Perrino, F.W., Hollis, T., 2007. The crystal structure of TREX1 explains the 3′ nucleotide specificity and reveals a polyproline II helix for protein partnering. Journal of Biological Chemistry 282, 10537–10543.

Dvir, H., Harel, M., Bon, S., Liu, W.Q., Vidal, M., Garbay, C., Sussman, J.L., Massoulie, J., Silman, I., 2004. The synaptic acetylcholinesterase tetramer assembles around a polyproline II helix. The EMBO journal 23, 4394–4405.

Hicks, J.M., Hsu, V.L., 2004. The extended left‐handed helix: A simple nucleic acid‐binding motif. Proteins: Structure, Function, and Bioinformatics 55, 330–338.

Hu, K.-N., Havlin, R.H., Yau, W.-M., Tycko, R., 2009. Quantitative determination of site-specific conformational distributions in an unfolded protein by solid-state nuclear magnetic resonance. J Mol Biol 3921, 1055–1073.

Hubbard, S.J., Thornton, J.M., 1993. Naccess. Computer Program, Department of Biochemistry and Molecular Biology, University College London 2.

Hung, L.-W., Kohmura, M., Ariyoshi, Y., Kim, S.-H., 1998. Structure of an enantiomeric protein, d-monellin at 1.8 Å resolution. Acta Crystallographica Section D: Biological Crystallography 54, 494–500.

Jha, A.K., Colubri, A., Zaman, M.H., Koide, S., Sosnick, T.R., Freed, K.F., 2005. Helix, sheet, and polyproline II frequencies and strong nearest neighbor effects in a restricted coil library. Biochemistry 44, 9691–9702.

Joshi, H.K., Etzkorn, C., Chatwell, L., Bitinaite, J., Horton, N.C., 2006. Alteration of sequence specificity of the type II restriction endonuclease HincII through an indirect readout mechanism. Journal of Biological Chemistry 281, 23852–23869.

Kay, B.K., Williamson, M.P., Sudol, M., 2000. The importance of being proline: the interaction of proline-rich motifs in signaling proteins with their cognate domains. The FASEB journal 14, 231–241.

Kelly, M.A., Chellgren, B.W., Rucker, A.L., Troutman, J.M., Fried, M.G., Miller, A.F., Creamer, T.P., 2001. Host-guest study of left-handed polyproline II helix formation. Biochemistry 40, 14376–14383.

Kieliszewski, M.J., Lamport, D.T., 1994. Extensin: repetitive motifs, functional sites, post-translational codes, and phylogeny. Plant J 5, 157–172.

Kim, C.-Y., Quarsten, H., Bergseng, E., Khosla, C., Sollid, L.M., 2004. Structural basis for HLA-DQ2-mediated presentation of gluten epitopes in celiac disease. Proceedings of the National Academy of Sciences 101, 4175–4179.

King, S.M., Johnson, W.C., 1999. Assigning secondary structure from protein coordinate data. Proteins 35, 313–320.

Kretsinger, R.H., Nockolds, C.E., 1973. Carp muscle calcium-binding protein. II. Structure determination and general description. J Biol Chem 248, 3313–3326.

Krimm, S., Tiffany, M.L., 1974. The circular dichroism spectrum and structure of unordered polypeptides and proteins. Israel Journal of Chemistry 12, 189–200.

Kumar, P., Bansal, M., 2012. HELANAL-Plus: a web server for analysis of helix geometry in protein structures. J Biomol Struct Dyn 30, 773–783.

Kumar, P., Bansal, M., 2015a. Dissecting pi-helices: sequence, structure and function. Febs J 282, 4415-4432.

Kumar, P., Bansal, M., 2015b. Identification of local variations within secondary structures of proteins. Acta Crystallogr D Biol Crystallogr 71, 1077–1086.

Kumar, P., Kailasam, S., Chakraborty, S., Bansal, M., 2014. MolBridge: a program for identifying nonbonded interactions in small molecules and biomolecular structures. Journal of Applied Crystallography 47, 1772–1776.

Kumar, S., Bansal, M., 1998. Dissecting alpha-helices: position-specific analysis of alpha-helices in globular proteins. Proteins 31, 460–476.

Larson, M.R., Rajashankar, K.R., Patel, M.H., Robinette, R.A., Crowley, P.J., Michalek, S., Brady, L.J., Deivanayagam, C., 2010. Elongated fibrillar structure of a streptococcal adhesin assembled by the high-affinity association of α-and PPII-helices. Proceedings of the National Academy of Sciences 107, 5983–5988.

Makarov, A., Lobachov, V., Adzhubei, I., Esipova, N., 1992. Natural polypeptides in left‐handed helical conformation A circular dichroism study of the linker histones’ C‐terminal fragments and β‐endorphin. FEBS Lett 306, 63–65.

Mansiaux, Y., Joseph, A.P., Gelly, J.C., de Brevern, A.G., 2011. Assignment of PolyProline II conformation and analysis of sequence-structure relationship. PLoS One 6, e18401.

MATLAB. 2010. version 7.10.0 (R2010a). The MathWorks Inc.

Mazik, M., Bläser, D., Boese, R., 2000. The potential of CH··· N interactions in determining the crystal structures of novel 3, 4-disubstituted-5-pyridinyl-isoxazoles. Tetrahedron Letters 41, 5827–5831.

Novotny, M., Kleywegt, G.J., 2005. A survey of left-handed helices in protein structures. J Mol Biol 347, 231–241.

Pauling, L., Corey, R.B., 1951. The pleated sheet, a new layer configuration of polypeptide chains. Proc Natl Acad Sci U S A 37, 251–256.

Pauling, L., Corey, R.B., Branson, H.R., 1951. The structure of proteins; two hydrogen-bonded helical configurations of the polypeptide chain. Proc Natl Acad Sci U S A 37, 205–211.

Pentelute, B.L., Gates, Z.P., Tereshko, V., Dashnau, J.L., Vanderkooi, J.M., Kossiakoff, A.A., Kent, S.B., 2008. X-ray structure of snow flea antifreeze protein determined by racemic crystallization of synthetic protein enantiomers. Journal of the American Chemical Society 130, 9695–9701.

Perutz, M.F., 1951. New x-ray evidence on the configuration of polypeptide chains. Nature 167, 1053-1054.

Pickering, A.L., Seeber, G., Long, D.-L., Cronin, L., 2005. The importance of pi-pi, pi-CH and N-CH interactions in the crystal packing of Schiff-base derivatives of cis, cis- and cis,trans-1,3,5-triaminocyclohexane. CrystEngComm 7, 504–510.

Punta, M., Coggill, P.C., Eberhardt, R.Y., Mistry, J., Tate, J., Boursnell, C., Pang, N., Forslund, K., Ceric, G., Clements, J., Heger, A., Holm, L., Sonnhammer, E.L., Eddy, S.R., Bateman, A., Finn, R.D., 2012. The Pfam protein families database. Nucleic Acids Res 40, D290–301.

Rucker, A.L., Creamer, T.P., 2002. Polyproline II helical structure in protein unfolded states: lysine peptides revisited. Protein Sci 11, 980–985.

Schrodinger, LLC. 2010. The PyMOL Molecular Graphics System, Version 1.3r1.

Shi, Z., Woody, R.W., Kallenbach, N.R., 2002. Is polyproline II a major backbone conformation in unfolded proteins? Adv Protein Chem 62, 163–240.

Shi, Z., Chen, K., Liu, Z., Kallenbach, N.R., 2006. Conformation of the backbone in unfolded proteins. Chemical Reviews 106, 1877–1897.

Siligardi, G., Drake, A.F., 1995. The importance of extended conformations and, in particular, the PII conformation for the molecular recognition of peptides. Biopolymers 37, 281–292.

Soman, K.V., Ramakrishnan, C., 1983. Occurrence of a single helix of the collagen type in globular proteins. J Mol Biol 170, 1045–1048.

Sreerama, N., Woody, R.W., 1999. Molecular dynamics simulations of polypeptide conformations in water: A comparison of α, β, and poly (pro) II conformations. Proteins: Structure, Function, and Bioinformatics 36, 400–406.

Srinivasan, R., Rose, G.D., 1999. A physical basis for protein secondary structure. Proc Natl Acad Sci U S A 96, 14258–14263.

Stapley, B.J., Creamer, T.P., 1999. A survey of left-handed polyproline II helices. Protein Sci 8, 587–595.

Syme, C.D., Blanch, E.W., Holt, C., Jakes, R., Goedert, M., Hecht, L., Barron, L.D., 2002. A Raman optical activity study of rheomorphism in caseins, synucleins and tau. European Journal of Biochemistry 269, 148–156.

Thauer, R.K., Kaster, A.-K., Seedorf, H., Buckel, W., Hedderich, R., 2008. Methanogenic archaea: ecologically relevant differences in energy conservation. Nat Rev Micro 6, 579–591.

Vila, J.A., Baldoni, H.A., Ripoll, D.R., Ghosh, A., Scheraga, H.A., 2004. Polyproline II helix conformation in a proline-rich environment: a theoretical study. Biophys J 86, 731–742.

Wang, G., Dunbrack, R.L., Jr., 2003. PISCES: a protein sequence culling server. Bioinformatics 19, 1589–1591.

Whittington, S.J., Chellgren, B.W., Hermann, V.M., Creamer, T.P., 2005. Urea promotes polyproline II helix formation: implications for protein denatured states. Biochemistry 44, 6269–6275.

Williamson, M.P., 1994. The structure and function of proline-rich regions in proteins. Biochemical journal 297, 249.

Wu, H., de Graaf, B., Mariani, C., Cheung, A.Y., 2001. Hydroxyproline-rich glycoproteins in plant reproductive tissues: structure, functions and regulation. Cell Mol Life Sci 58, 1418–1429.

Zarrinpar, A., Bhattacharyya, R.P., Lim, W.A., 2003. The structure and function of proline recognition domains. Homo 332, 20.

Zawadzke, L.E., Berg, J.M., 1993. The structure of a centrosymmetric protein crystal. Proteins: Structure, Function, and Bioinformatics 16, 301–305.

